# HiCMC: High-Efficiency Contact Matrix Compressor

**DOI:** 10.1101/2023.11.03.565487

**Authors:** Yeremia Gunawan Adhisantoso, Tim Körner, Fabian Müntefering, Jörn Ostermann, Jan Voges

## Abstract

Chromosome organization plays an important role in biological processes such as replication, regulation, and transcription. One way to study the relationship between chromosome structure and its biological functions is through Hi-C studies, a genome-wide method for capturing chromosome conformations. Such studies generate vast amounts of data. The problem is exacerbated by the fact that chromosome organization is dynamic, requiring snapshots at different points in time, further increasing the amount of data to be stored. We present a novel approach called the High-Efficiency Contact Matrix Compressor (HiCMC) for efficient compression of Hi-C data. By modeling the underlying structures found in the contact matrix, such as compartments and domains, HiCMC outperforms CMC by approximately 8% and more than 50% against cooler, LZMA, and bzip2 over the state of the art across multiple cell lines and resolutions. In addition, the domain information that is embedded in the data can be used to speed up downstream analysis. HiCMC is available at https://github.com/sXperfect/hicmc.

## 1. Introduction

The human genome provides critical insights into a wide range of biological processes. In recent decades, advances in high-throughput sequencing technologies have reduced the costs associated with genome sequencing [32]. This cost reduction has enabled large-scale studies such as genome-wide association studies [25] and the development of the concept of polygenic risk scores [12]. These studies involve the systematic analysis of hundreds of thousands of genetic variants associated with specific traits or diseases. They unravel many complex interactions between genotypes and phenotypes.

Simultaneously, the advances in high-throughput sequencing technologies have spurred advances in the field of epigenetics [13], i.e., the study of biological processes that do not involve alterations directly in the underlying DNA sequence, but with regard to other genetic features such as spatial organization of chromosomes. One of the most important findings has been indeed the critical role of chromosome spatial structure in biological functions such as replication, regulation, and transcription [7, 30]. One way to analyze the three-dimensional structure of the chromosome is through chromosome conformation capture (3C) [10], a ligation-based approach that captures the interactions between pairs of loci. 3C successors such as Hi-C and Micro-C [17, 22, 27, 33] are able to capture genome-wide interactions between all possible pairs of loci of all chromosomes simultaneously and with much higher resolution. Hi-C and Micro-C allow the identification of long-range interactions and provide insight into finer chromosomal structures such as topologically associating domains (TADs) and loop domains [9,28]. Figure 1 shows an example of an so-called intra-chromosomal (*cis*) contact matrix as a result of a Hi-C experiment. In the figure, highly-interacting regions are colored in dark red while regions with a low amount of interactions are colored in brighter shades of red. From the figure, it can hence be seen, e.g., by the dark red diagonal, that interactions are highly correlated with spatial proximity. Each row and column of the contact matrix represents a locus of a given size. The size of the loci is defined as the resolution. A high resolution contact matrix means that the contact matrix can reveal finer structure, therefore the size of the loci is small.

**Fig. 1:**
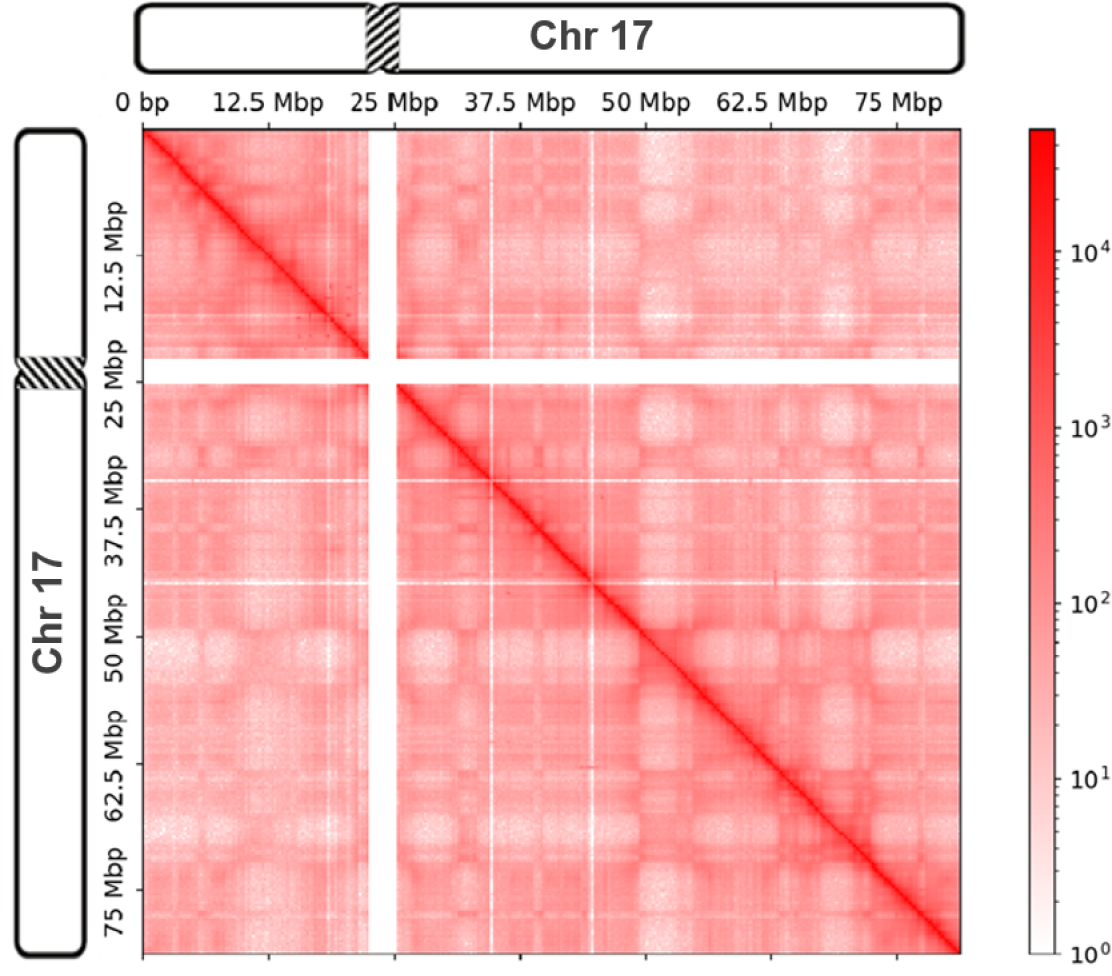
An example of an intra-chromosomal (*cis*) contact matrix of chromosome 17. Interactions are highly correlated with spatial proximity, and hence in the figure, highly-interacting regions are colored in dark red while regions with a low amount of interactions are colored in brighter shades of red. The contact matrix is sparse, symmetrical and contains regions with no interactions, shown as white rows and columns.

Such experiments generate enormous amounts of data, in particular when they are performed with a highresolution, i.e., interactions are counted at a small granularity. In addition, the three-dimensional organization of chromosomes is dynamic. It changes over time and exhibits cell type specificity. Thus, comprehensive analyses require the examination of chromosome organization across multiple temporal snapshots, compounding the challenge of data volume. While there are established standards for representing DNA sequencing data, such as the FASTQ [6], SAM [21], and MPEG-G [34] formats, a common format or method for representing and compressing three-dimensional interactions that can accommodate large data sets is lacking [8].

Several formats have been developed to provide efficient storage of Hi-C data, such as hic [14] and butlr [35]. Later, cooler [2], based on the existing HDF5 [20] format, was introduced. The HDF5 format provides flexible organization of multidimensional arrays, support for random access, and data compression based on Zlib [11] and sZIP [37]. Cooler takes advantage of sparsity and symmetry properties by storing contact matrices in Coordinate List (COO) representation. However, the performance of HDF5 compression is inferior compared to modern general-purpose compression methods such as Lempel–Ziv–Markov chain algorithm (LZMA) [26] and bzip2 [29], and it does not exploit the chromosomal structures found in the contact matrix. In contrast to the aforementioned formats, Contact Matrix Compressor (CMC) [3] improves compression performance by exploiting several properties of the contact matrix, including the correlations between genomic distance and interactions, unalignable regions, and symmetry. While it improves compression, it does not take advantage of the finer structures found in the contact matrix, such as compartments and TADs. In this work, we present a novel approach, High-Efficiency Contact Matrix Compressor (HiCMC), for the contact matrix compression. Better performance is achieved by modeling hierarchical structures in the chromosomes, such as TADs.

## 2. Methods

Our approach HiCMC is a major extension of CMC [3]. It comprises splitting the genome-wide contact matrix into intra- and inter-chromosomal sub-contact matrices, row and column masking, model-based transformation, row binarization, and entropy coding as shown in Figure 2. The key idea of CMC is to sort the matrix values such that in each row of a contact matrix, the number of bits required for each value, i.e., the magnitude of the values, is similar. Lieberman-Aiden et al. [22] observed that the probability of contact can be viewed as a function of distance for contacts within a chromosome. The main drawback of CMC is that it does not account for the fact that chromosomes create structures such as compartments and domains that are highly interacting with themselves. Such structures cause the interactions in certain regions of the contact matrix to be lower or higher than the expected interactions based on the distance. Therefore, we sort the values based on the prediction of a model. We will discuss these processes in detail in the following sections.

**Fig. 2:**
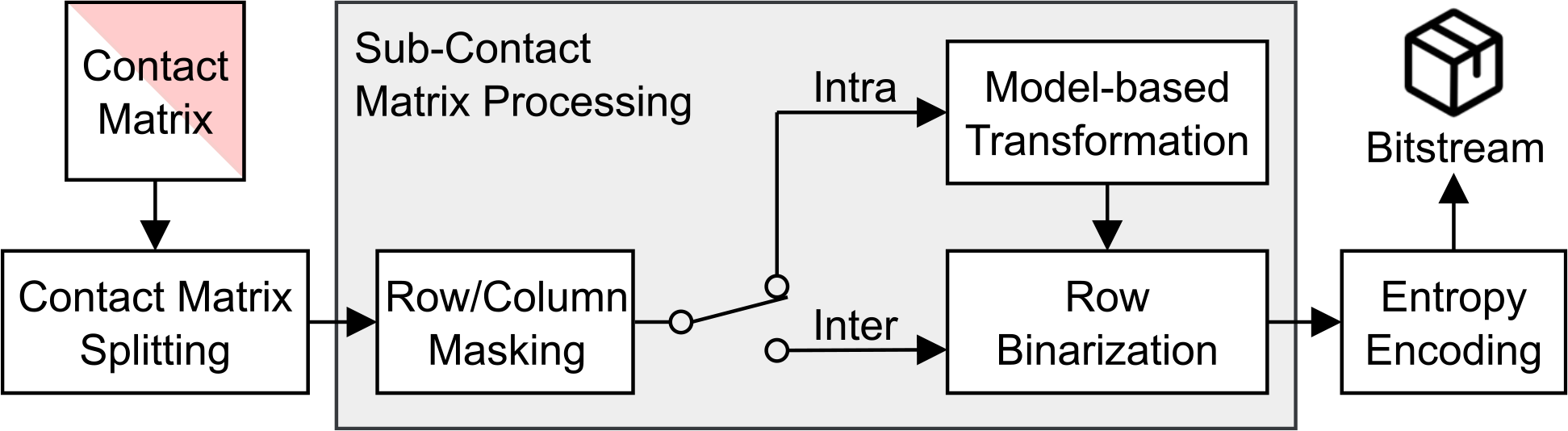
The HiCMC compression pipeline consists of splitting the genome-wide contact matrix into intra- and inter-chromosomal contact matrices, row/column masking, model-based transformation, row binarization, and entropy coding. The type of input sub-contact matrix determines whether Intra or Inter is used.

### 2.1 Split Contact matrix

The first step in the HiCMC compression pipeline is to divide the contact matrix into chromosome-chromosome interaction matrices, hereafter referred to as sub-contact matrices. Due to the symmetry of contact matrices, only sub-contact matrices lying within the upper triangle need to be stored. The contact matrix after splitting is shown in the Figure 3a. The pipeline is branched differently depending of the type of sub-contact matrix. Additional transformations are done to the intra-chromosomal sub-contact matrices because they are denser and contain more structures than the inter-chromosomal sub-contact matrices. In addition, chromosomal structures are found in the intra-chromosomal sub-contact matrix rather than the inter-chromosomal sub-contact matrix. By splitting the genome-wide contact matrix, we can differentiate the pipeline based on the statistics we want to exploit.

**Fig. 3:**
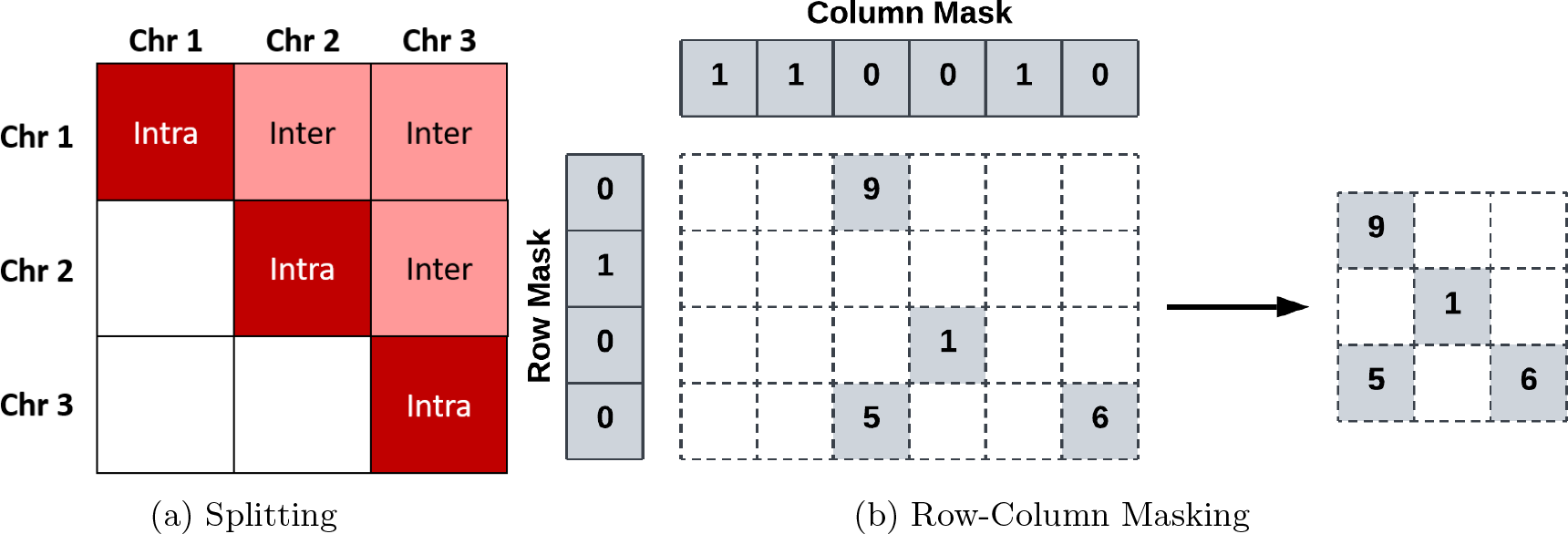
Splitting and masking processes of HiCMC. (a) The contact matrix is divided into two different sub-contact matrices based on chromosome-chromosome interactions: intra-chromosomal and inter-chromosomal sub-contact matrix. We only store the sub-matrices that lie in the main and upper triangle of the matrix. (b) The masking process works by marking empty rows/columns in the corresponding mask (left) and then removing them from the original matrix to construct the masked matrix (right).

### 2.2 Row-Column Masking

To remove redundant information in sub-contact matrices efficiently, we next remove unalignable regions [3] – rows or/and columns with no interactions – by first marking the rows and columns with the binary masks (see Figure 3b). The mask entry is set to 1 for the corresponding rows and columns that exclusively contain zeros. Conversely, in cases where there is at least one non-zero entry, the mask entry is set to 0. The marked rows and columns are subsequently eliminated from the sub-contact matrix.

### 2.3 Model-based Transformation

The diagonal transformation of CMC assumes that the values in a diagonal of the contact matrix are of approximately similar magnitude This transformation reflects the observation that the chromosomal interactions serve as an approximation of spatial distance [22]. By placing the entries from the same diagonal in a row in the new matrix, the number of bits required to represent the values in each row can be reduced, as shown in Figure 4. However, due to lower-level features such as A/B compartments or TADs it provides only a basic approximation, as the average interactions between compartments or domains may vary. Interactions within TAD are enriched, but an abrupt drop in interactions is observed for inter-TADs [27].

**Fig. 4:**
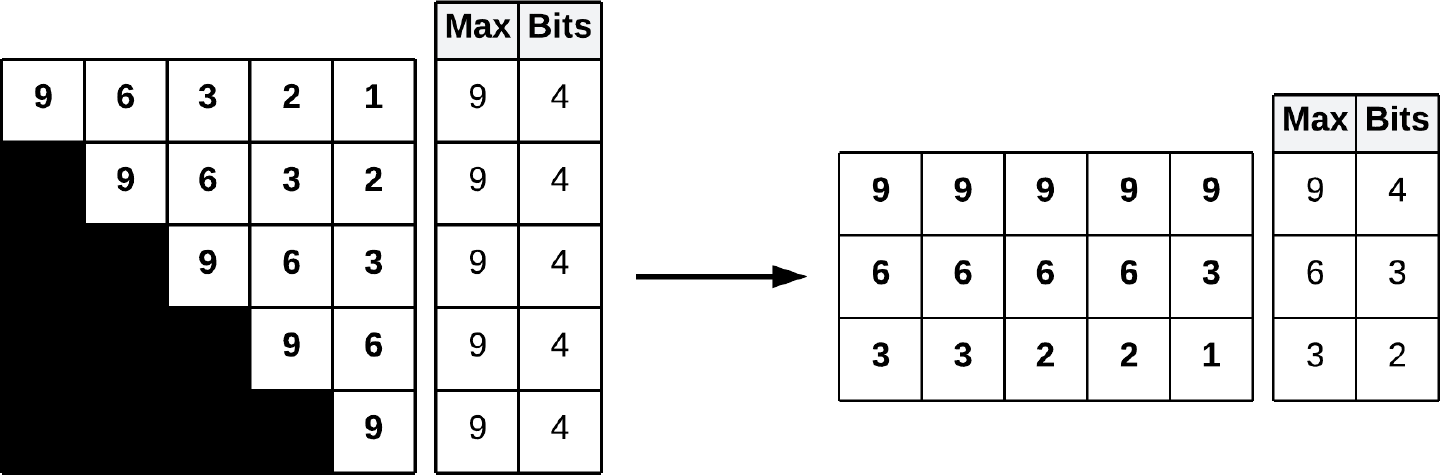
The diagonal transformation of CMC iteratively places entries from each diagonal, starting from the main diagonal, into the output matrix (right), resulting in fewer bits needed to represent each value.

The model-based transformation approach is designed to overcome these limitations. Model-based transformation consists of 3 main steps, as shown in Figure 5. First, we predict the entries of the contact matrix based on our model. Preferably, this model should take into account the correlations introduced by lower-level features such as A/B compartments and TADs to more accurately capture the underlying structure of the contact matrix. We then determine the sorting indices by sorting the entries of the contact matrix by their magnitude. The indices are then used to sort the contact matrix by placing each entry of the original matrix in its corresponding index.

**Fig. 5:**
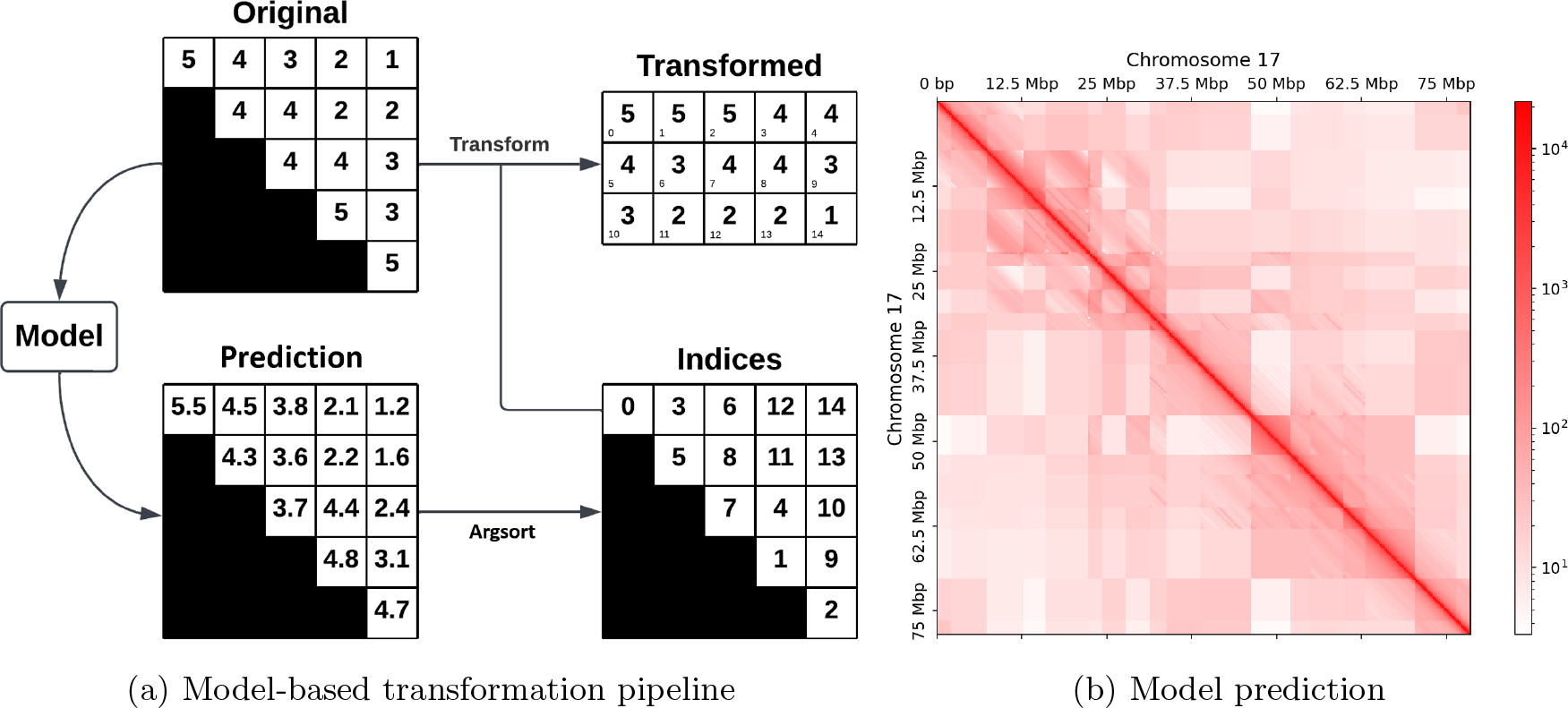
Overview of the model-based transformation pipeline and model prediction. (a) First, the modelbased transformation predicts the interactions for each possible pair based on a model. Then, the prediction is sorted by its magnitude, yielding sorting indices (argsort operation). The original interactions are then placed in a new matrix based on these indices, starting from the top left to the bottom right. (b) An example of prediction generated by our model derived from Figure 1. Domain matrices modeled using the genomic distance function are distinguishable from matrices modeled using a constant domain value.

Prior to modeling, we determine the domain boundaries using TAD-caller and divide the sub-contact matrix into rectangular regions, hereafter called domain matrices. In our pipeline, we predict the domains based on the insulation score [15]. A key consideration when building the model is that the model must be included in the compressed payload, and therefore introduces an overhead. This represents a trade-off between the quality of our model and the compression performance. Hence, it is important to explore different methods to construct and efficiently encode these models. Finally, due to biases such as GC-rich regions and low mapability regions, the visibility across regions in a chromosome is affected. To further improve model accuracy, the model is constructed from a balanced matrix—using a method such as Knight-Ruiz normalization (KR)—thus removing the experimental bias introduced in the experiments.

The model approximates the entries of each domain matrices by either a constant value called domain value or a function of genomic distance as shown in Figure 5b. This is beneficial because the average interactions are roughly constant and no longer correlate with the genomic distance when the genomic distance is greater than a certain value or for inter-domain interactions [22, 27]. The decision for modeling each domain matrix depends on the statistical characteristic of the the domain matrix entries such as mean, sparsity, or standard deviation, and the corresponding threshold. Both statistical characteristic and the corresponding threshold are encoding parameters. The decision is stored as a binary matrix called domain mask with each entry represents the decision for each domain.

For domain construction using a function of genomic distance, we iterate over all domains and over all domain matrices, replacing each entry of the current domain matrix with the domain-wide average for entries at the same genomic distance. These values are stored in an auxiliary matrix called the distance table, which is constructed incrementally as we iterate over the domain matrices. Each column of the distance table stores all values for a given genomic distance between two loci. After calculating the average interactions for given genomic distances in a domain matrix, the average interaction values are added to the distance table in their corresponding columns. By constructing the distance table in this way, values of similar magnitude are placed in the same column, which can be exploited during entropy coding. This process can be seen in Figure 6.

**Fig. 6:**
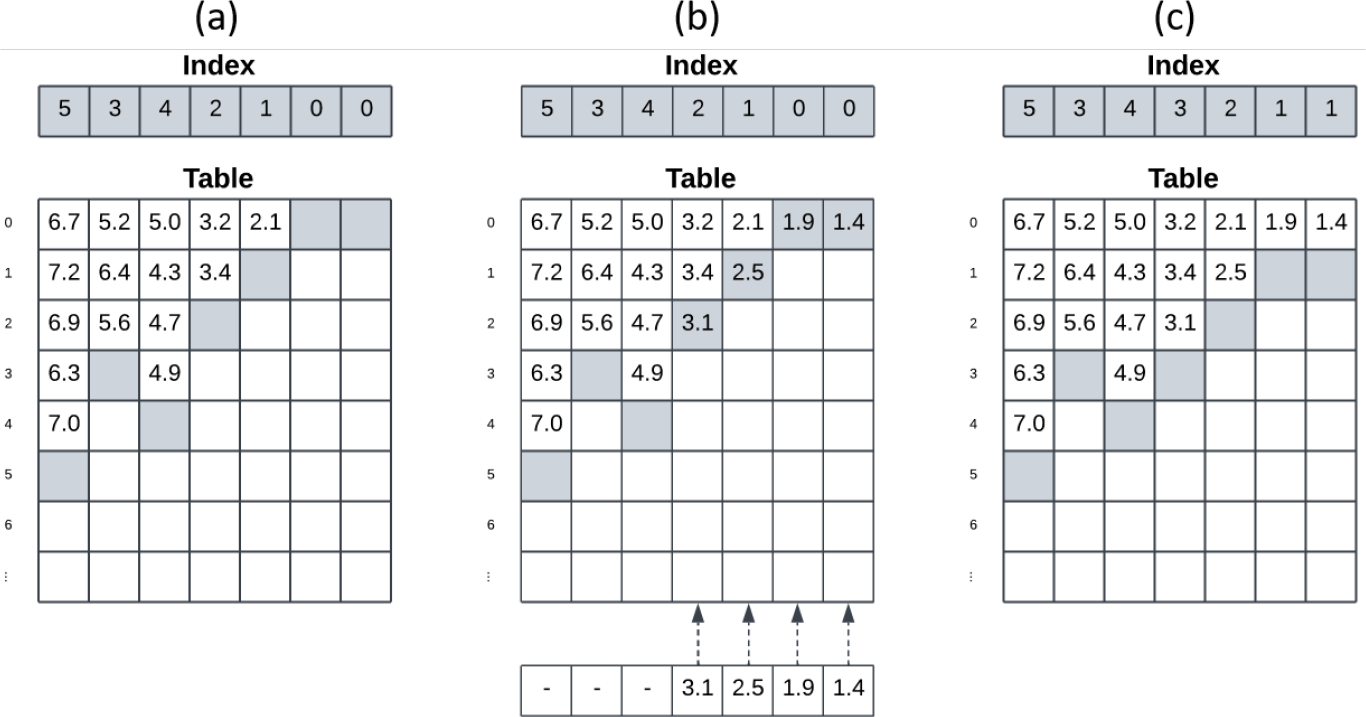
The process of inserting new values into an existing distance table. (a) The distance table consists of two parts: the index array and the table that keeps track of the interactions. Each column of the distance table stores all values for a given genomic distance between two loci, and the index array keeps track of the number of interactions stored in each column. (b) The values are added to the columns in the corresponding row indices. (c) The entries in the index array are incremented for the columns processed.

**Fig. 7:**
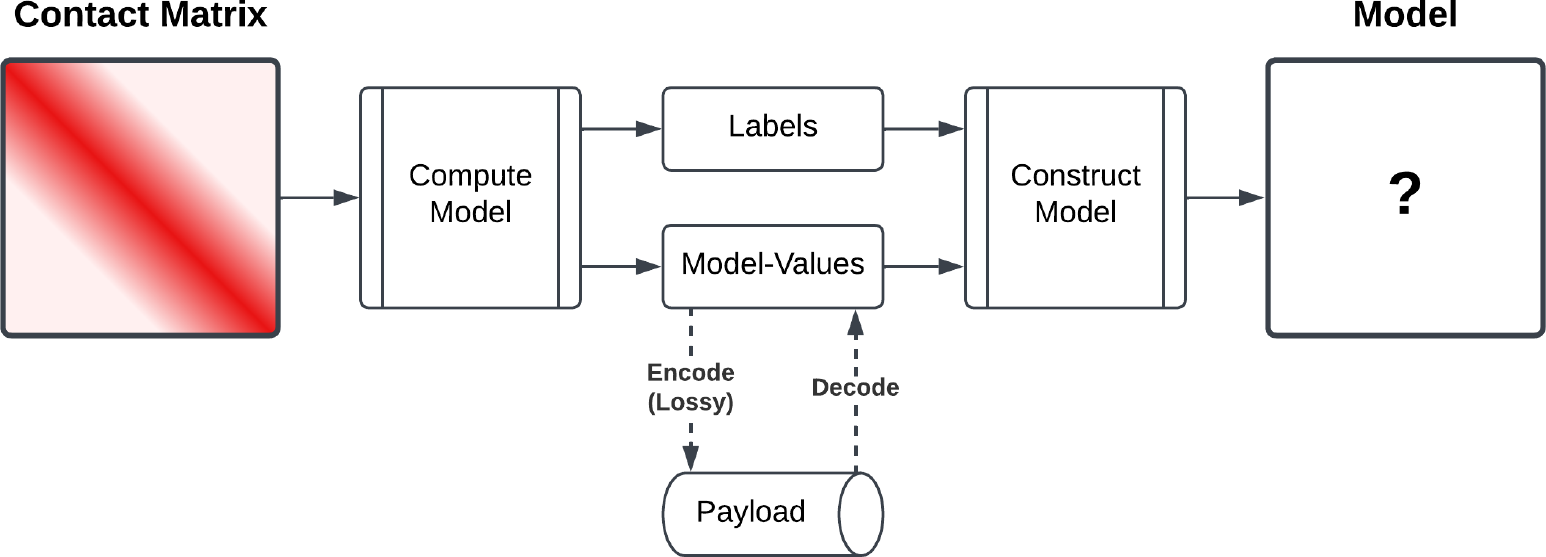
The model can be reconstructed using two payloads: the domain labels and the model values. While the domain labels are encoded lossless, the model values can optimally be compressed using a lossy encoding method.

For domain construction using a domain value, the domain matrix is modeled using the average interactions of the current domain matrix. These additional values can be efficiently stored in another auxiliary structure, a two-dimensional matrix that stores the contact average for each domain matrix, which hereafter referred to as domain values. An example of a predicted model is shown in Figure 5b.

### 2.4 Row Binarization

After the model-based transformation is applied, the values in each row are approximately similar in magnitude (see Figure 5a). In the row binarization transformation [3], we decompose each row into binary rows, resulting in binary rows starting from the binary row that represents the least significant bit up to the binary row that represents the most significant bit. The bit length of *i*-th row *q*_*i*_ depends on the greatest value in a row and is computed as

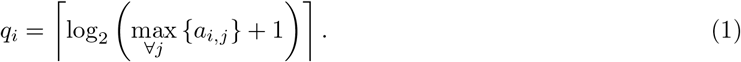

where *a*_*i,j*_ is the value of the transformed matrix at row *i* and column *j*. Instead of storing the bit length separately, this value is represented as the so-called sentinel value by prepending the sentinel value to the matrix as the first column.

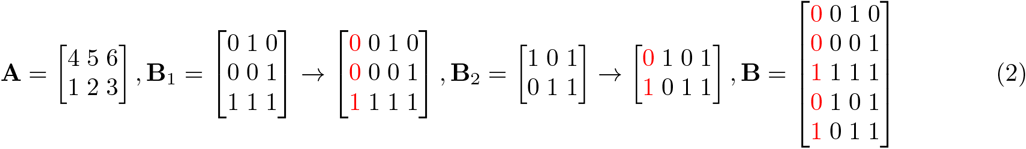

To illustrate the steps, we represent them in Equation 2: Suppose we have a transformed matrix **A**. The first and second rows of **A** are decomposed into binary rows, yielding binary matrices **B**_1_ and **B**_2_, respectively. The first row of these matrices represents the least significant bits of the original value, and the last row represents the most significant bits. For each binary matrix **B**_*i*_, an additional column is prepended to the matrix to indicate whether a particular binary row represents the most significant bit. Value 1 for the most significant bit, otherwise 0. Finally, both matrices **B**_1_ and **B**_2_ are concatenated into a single binary matrix **B**.

### 2.5 Entropy Coding

In total, four payloads are required for the model: the TAD boundaries, the domain mask, the domain values, and the distance table. The TAD boundaries can be represented as a one-dimensional binary array indicating the presence or absence of a boundary for each bin. It can be efficiently encoded using binary run-length encoding [3], since long sequences of zeros (indicating the absence of a boundary) are expected.

The domain mask is a two-dimensional, square, symmetric, binary matrix. It indicates for each domain matrix whether the interactions are modeled as a function of genomic distance or by a simple average. Since there are many 1’s along the main diagonal of the matrix, it is first transformed using the diagonal transformation and then compressed using encoder compliant to the JBIG standard (ISO/IEC 11544 [18]), specialized for lossless compression of bi-level (binary) images. It takes advantage of the spatial correlation of binary pixels or contexts (in this case, values of neighboring pixels). The length of the context varies between a total of 10 to 12 neighboring values from the previous rows and columns, depending on the context mode. The domain values have a form similar to a domain mask. The entries are the mean of the corresponding domain matrix. It is also transformed using the diagonal transformation, as higher values tend to be placed along the main diagonal. The matrix is then compressed using fpzip [23], a compression algorithm used for lossless or lossy compression of large multi-dimensional floating-point arrays that exhibit spatial correlation. Similarly, the distance table matrix is encoded by serializing it into an array, which is also compressed using fpzip. Finally, the transformed interactions of the contact matrix are compressed using prediction by partial patching (PPM) [16]-based technique PPMd [31]. We use the lossy mode of fpzip to compress all floating-point values with floating-point precision as the parameter.

## 3 Results

For the evaluation, we use the dataset published by Rao et al. [27] and available under the NCBI accession code GSE63525. The dataset consists of contact matrices from various cell lines (from the species *H. sapiens* and *M. musculus*) at multiple resolutions. We convert the data into the cooler format using hic2cool tool. The dataset is also available on the 4D Nucleome Project data portal. The dataset is described in Table 1. Our method, HiCMC, is available at https://github.com/sXperfect/hicmc.

**Table 1:** The dataset used for evaluation consists of contact matrices at multiple resolutions from different cell lines and based on different approaches. Cell line hic [GB] cooler [GB] cooler (intra) [GB].

| Cell line | hic [GB] | cooler [GB] | cooler (intra) [GB] |
| --- | --- | --- | --- |
| CH12 | 8.58 | 1.90 | 0.43 |
| GM12878 (Insitu-DpnII) | 6.83 | 1.37 | 0.44 |
| GM12878 (Insitu-Primary) | 31.86 | 9.64 | 1.77 |
| GM12878 (Insitu-Replicate) | 29.06 | 8.78 | 1.63 |
| HMEC | 7.08 | 1.46 | 0.20 |
| HUVEC | 8.87 | 1.92 | 0.27 |
| IMR90 | 12.77 | 2.72 | 0.44 |
| K562 | 12.08 | 2.61 | 0.37 |
| KBM7 | 13.91 | 3.19 | 0.33 |
| NHEK | 11.34 | 2.55 | 0.62 |

Since our approach extends the compression pipeline of CMC, we limit the comparison to the intrachromosomal contact matrices. As a pre-processing step before the actual compression, we predict the domain boundaries for each intra-chromosomal contact matrix using TAD-callers based on the insulation score [15] that is integral component of cooltools [1]. The contact matrices are balanced using KR [19] algorithm. The compression process in HiCMC is controlled by five encoding parameters: window size, threshold, distance table precision, domain value precision, and balancing weight precision. The domain border is determined by the insulation score, which aggregates interactions in a sliding window along the diagonal. The Insulation score has a window size parameter that specifies the size of the previously mentioned sliding window. The domain table precision, domain values precision, and balancing weights precision specify the floating-point precision for encoding of the corresponding payloads using fpzip. Last, the threshold determines the threshold value used to select a mode for the domain: representing a domain with its average or as a function of genomic distance. The parameters are optimized using the Tree-structured Parzen Estimator (TPE) algorithm [4, 5]. The resolution-specific parameter sets are described in Table 2. For compression using LZMA and bzip, the contact matrices undergo conversion into GInteractions [24] format using the HiCExplorer [36] tool. Subsequently, the matrices are compressed using their corresponding software and default parameters.

**Table 2:**
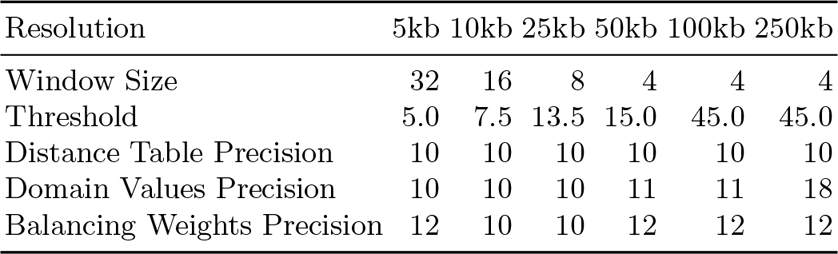
The resolution-specific parameter sets used by our compression pipeline. The parameter values are optimized using TPE algorithm.

As shown in Figure 8, HiCMC outperforms all other methods in terms of compression for intra-chromosomal contact matrices across all resolutions and cell lines. HiCMC performs better than CMC by around 8% on average for encoding of intra-chromosomal contact matrices and, more significantly, by over 50% compared to cooler, LZMA, and bzip2.

**Fig. 8:**
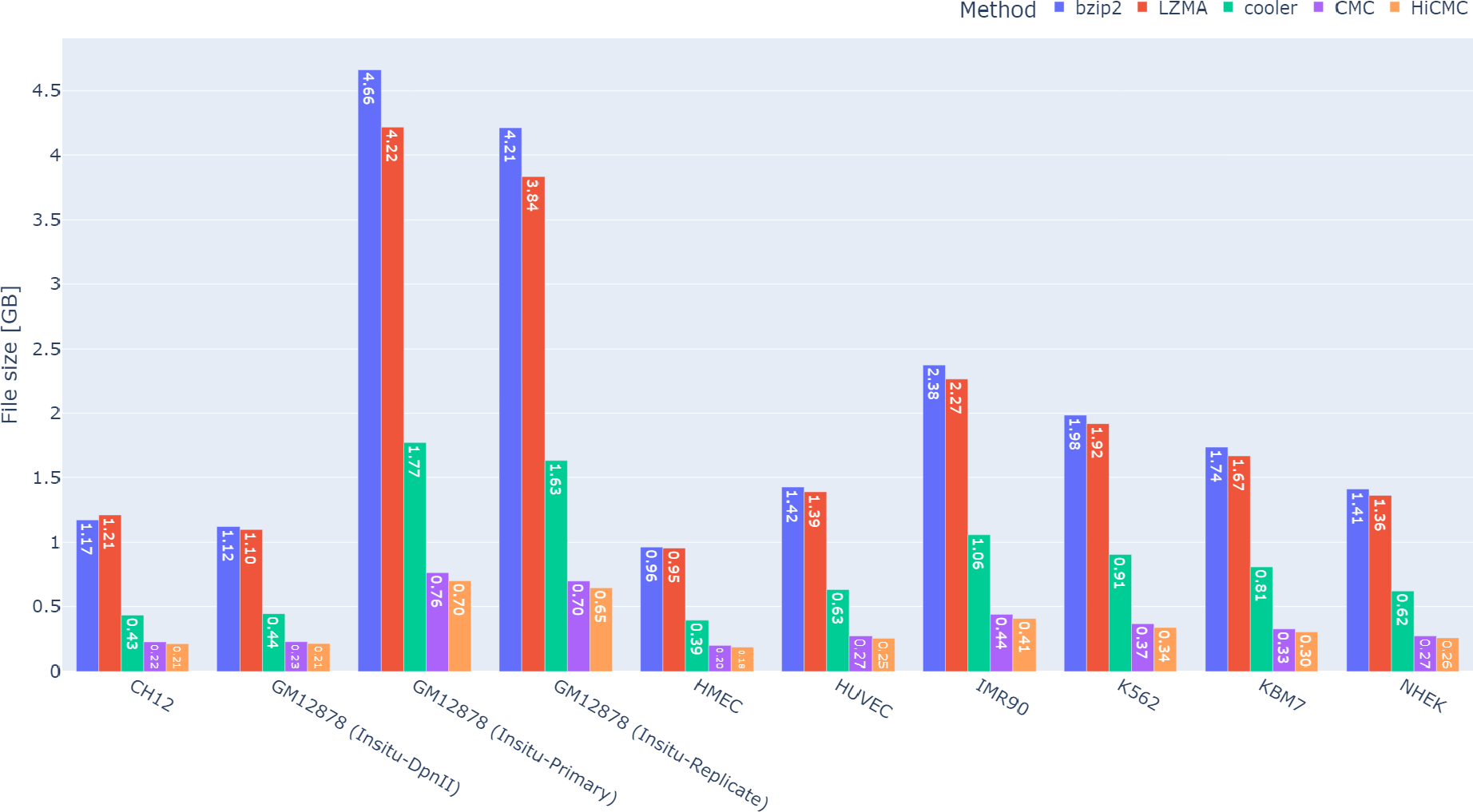
Absolute payload sizes of the compressed intra-chromosomal contact matrices. HiCMC outperforms CMC, cooler, LZMA, and bzip2 across all resolutions and cell lines.

To be more specific, we compare HiCMC to CMC for each resolution across all cell lines shown in Figure 9. Because the size of the domains is relatively large, it is most efficient to compress data at medium resolution (25 kb to 100 kb). The compression of the contact matrix can be further improved by experimenting with other TAD-callers, as we mentioned earlier in Section 2.3. We found that for higher resolution data (10kb and above) the contact matrix are very sparse, greater than 90%, as shown in Figure 10. A significant portion of the compressed payload is allocated to store the coordinates of observed interactions rather than actual interaction data. To further enhance compression performance for higher resolution contact matrices, we believe that better sparse binary matrix compression methods would be beneficial to improve the compression performance.

**Fig. 9:**
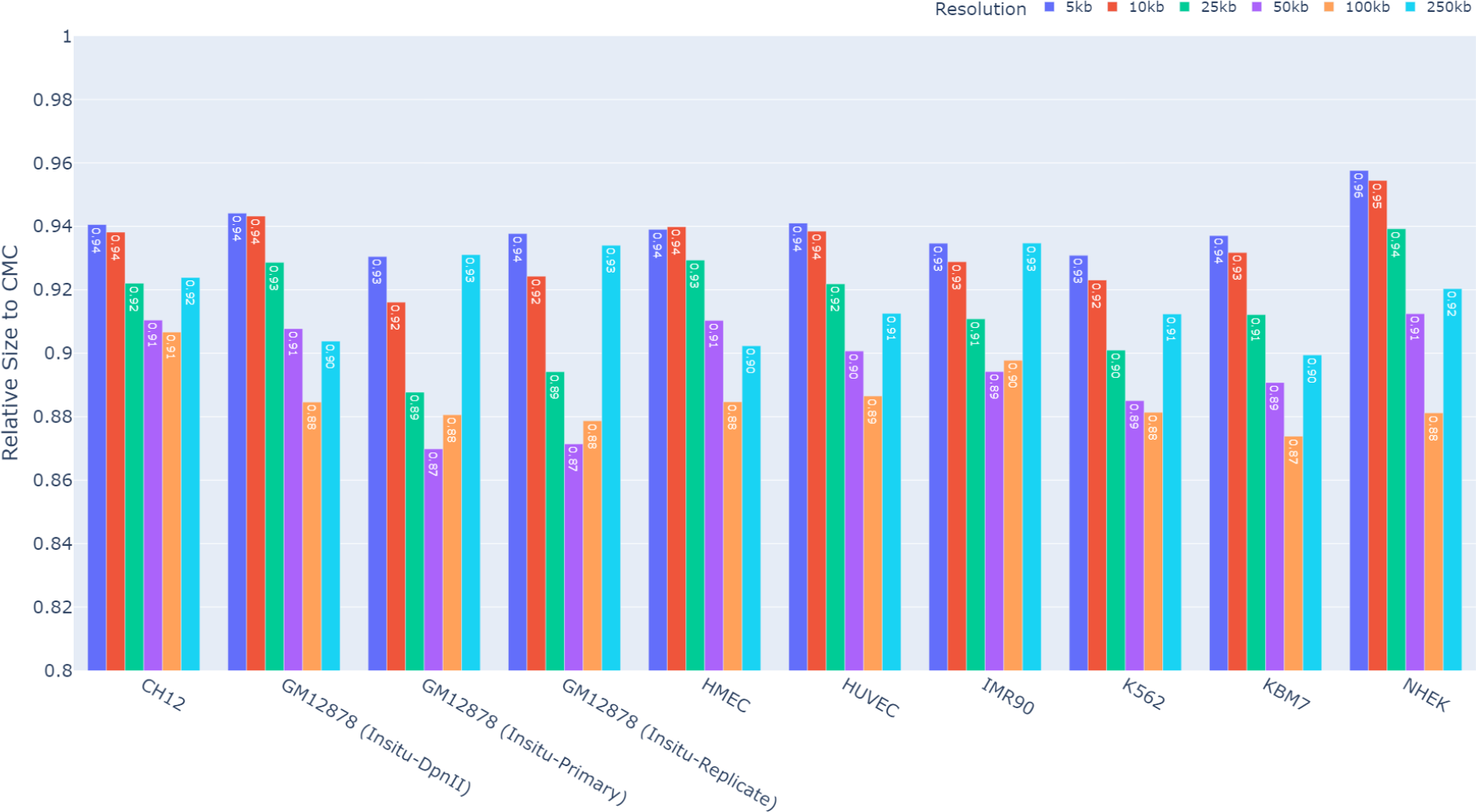
Relative size of the compressed HiCMC payload in comparison to that of CMC. HiCMC outperforms CMC across all resolutions and cell lines.

**Fig. 10:**
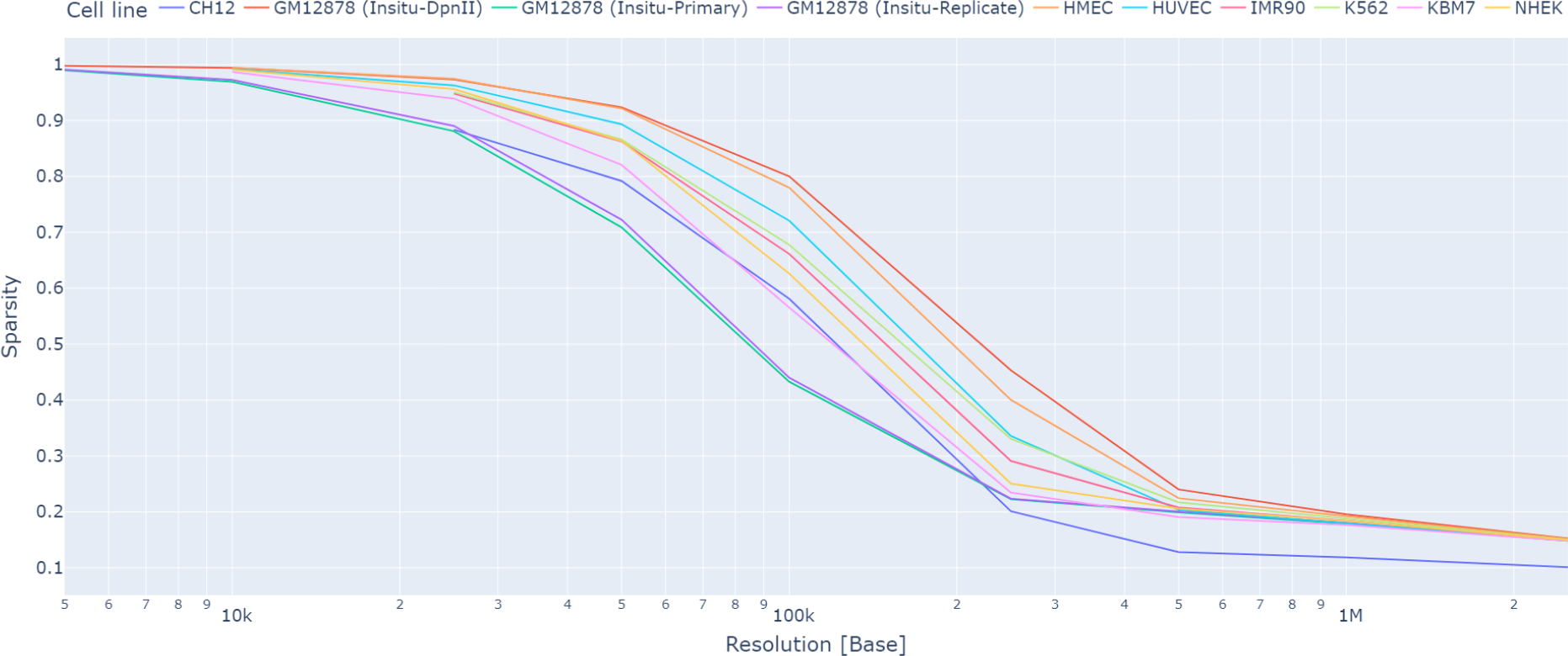
The sparsity of the contact matrix is a function of the resolution of the matrix. When the resolution is higher than 20 kb (lower value), the sparsity is greater than 90%. We suggest that improved sparse matrix transformation and encoding methods would be advantageous for encoding high-resolution contact matrices.

## 4 Conclusions

We have presented HiCMC, a specialized model-based compressor for encoding contact matrices from 3Cbased experiments. It outperforms cooler, generic compressors LZMA and bzip2, as well as the specialized contact matrix compressor CMC. HiCMC outperforms CMC by approximately 8% and is superior to other approaches for encoding of intra-chromosomal contact matrices by a minimum of 50%. HiCMC achieves better performance by exploiting the underlying properties of the contact matrix, such as symmetric, diagonally dominant, and hierarchical structures of chromosomal organization, in particular TAD. To exploit the hierarchical structure in the chromosome, the domains are modeled and used to predict the values of the contact matrix. For domain modeling, the boundaries of the domains are determined based on the insulation score. The model is adaptively determined based on the normalized value of the contact matrix to remove bias in the data. Furthermore, the domain information that is embedded in the data can be used to speed up downstream analyses.

## 5 Acknowledgements

The authors acknowledge the financial support by German Federal Ministry of Education and Research (BMBF) in the framework of personalisierte, prädiktive, präzise und präventive Medizin zur Verbesserung der Früherkennung, Diagnostik, Therapie und Prävention depressiver Erkrankungen (P4D) project under project number 01EK2204F. Views expressed herein are solely those of the author(s) and do not necessarily reflect the views of the German Federal Government nor the granting authority.

